# EEG-based age-prediction models as stable and heritable indicators of brain maturational level in children and adolescents

**DOI:** 10.1101/407049

**Authors:** Marjolein M.L.J.Z. Vandenbosch, Dennis van’t Ent, Dorret I. Boomsma, Andrey P. Anokhin, Dirk J.A. Smit

## Abstract

The human brain shows remarkable development of functional brain activity from childhood to adolescence. Here, we investigated whether electroencephalogram (EEG) recordings are suitable for predicting the age of children and adolescents. Moreover, we investigated whether over-or underestimation of age was stable over longer time periods, as stable prediction error can be interpreted as reflecting individual brain maturational level. Finally, we established whether the age-prediction error was genetically determined. Three minutes eyes-closed resting state EEG data from the longitudinal EEG studies of Netherlands Twin Register (n=836) and Washington University in St. Louis (n = 702) were used at ages 5, 7, 12, 14, 16 and 18. Longitudinal data were available within childhood and adolescence. We calculated power in 1 Hz wide bins (1 to 24 Hz). Random Forest regression and Relevance Vector Machine with 6-fold cross-validation were applied. The best mean absolute prediction error was obtained with Random Forest (1.22 years). Classification of childhood vs. adolescence reached over 94% accuracy. Prediction errors were moderately to highly stable over periods of 1.5 to 2.1 years (0.53 < r < 0.74) and signifcantly affected by genetic factors (heritability between 42% and 79%). Our results show that age prediction from low-cost EEG recordings is comparable in accuracy to those obtained with MRI. Children and adolescents showed stable over- or underestimation of their age, which means that some participants have stable brain activity patterns that reflect those of an older or younger age, and could therefore reflect individual brain maturational level. This prediction error is heritable, suggesting that genes underlie maturational level of functional brain activity. We propose that age prediction based on EEG recordings can be used for tracking neurodevelopment in typically developing children, in preterm children, and in children with neurodevelopmental disorders.

---

The neural tissue of the brain shows remarkable development from childhood to adolescence, which includes changes in dendritic arborization, synaptogenesis, and myelination, and synaptic pruning (Anderson, Northam, Hendy, & Wrennall, 2001; Huttenlocher, 1979). These neuronal-level processes result in brain volume increases and grey matter changes (Giedd et al., 2009;) Hedman, van Haren, Schnack, Kahn, & Hulshoff Pol, 2012; (Giedd et al., 2009; Hedman et al., 2012; Lenroot & Giedd, 2006; Mills & Tamnes, 2014a). These anatomical changes are accompanied by changes in brain function as reflected in electrophysiological brain activity. One of the most striking features is the change in oscillatory patterns in the electroencephalogram (EEG) (Niedermeyer & Lopes Da Silva, 2005; Smit et al., 2012). During childhood, temporal and posterior theta rhythm (4–7 Hz) dominates (Puligheddu et al., 2005 (Benniger, Matthis, & Scheffner, 1984)), but strongly decreases over the years. The alpha rhythm increases in frequency from 8 Hz in childhood to 10 Hz in adolescence (Smit et al., 2012). Maturation of alpha rhythms begins in posterior regions and ends in anterior regions while the beta frequency (12-30 Hz) matures from central to lateral and finally to frontal regions (Gasser, Jennen-Steinmetz, Sroka, Verleger, & Mocks, 1988; Niedermeyer & Lopes Da Silva, 2005). (Matousek & Petersen, 1973; Bresnahan, Anderson, & Barry, 1999) (Barriga-Paulino, Flores, & Gomez, 2011; Benniger et al., 1984; Gasser, Verleger, Bächer, & Sroka, 1988).

During maturation, children and adolescents show marked development of behavioral skills and cognition (Mills & Tamnes, 2014; Walhovd, Tamnes, & Fjell, 2014). Interestingly, they also show large differences in developmental speed. (Fischer & Silvern, 1985; Shaw et al., 2006). One of the challenges of neurodevelopmental research is to investigate how these differences in behavioral development can be explained by underlying changes in neural function (Durston, Casey, Durston, & Casey, 2006). Several studies have attempted to create measures of brain maturation by using brain imaging data for predicting calendar age—often using machine learning. These studies have largely focused on brain anatomy derived from magnetic resonance imaging (MRI). For example, Franke et al., (2012) computed the so-called *brain age* in children and adolescents and obtained an average prediction error of 1.1 years (defined as the average absolute difference between the subjects’ ages and the ages estimated by the machine learning model. Brown et al., (2012) predicted calendar age using the multidimensional nature of brain anatomy; At age 3 they obtained an average prediction error of 0.66 years. The prediction error increased with age until an average of 1.42 years at age 20.

These studies shows that anatomical brain maturation in childhood and adolescence can be estimated using expensive MRI scanning, which may result in limited availability. By contrast, EEG is inexpensive, and EEG units can be flexibly deployed in the field including educational and medical institutions. Moreover, the brain activity measured with EEG is known to show large developmental changes within the critical developmental periods of school-aged children (e.g., Smit et al., 2012). Our aim is to investigate whether EEG can be used to estimate the participants’ age with the same level of accuracy as obtained with MRI. To this end, we applied machine learning with cross-validation to predict age from resting-state EEG in a large sample of children and adolescents (5-18 years).

The previous studies also have shown that predicting calendar age is not perfect, and always results in (sometimes substantial) residual error. It has been suggested that this error is a biomarker of brain maturation (or *brain age*, Franke et al., 2012), since it indicates that some participants have brain patterns that are more appropriate for a different age than their own. Moreover, it has been suggested that the estimated brain maturation reflects the behavioral changes observed at an individual level, or correlates with neurodevelopmental disorders. However, this is arguably only the case if the prediction error is stable, and is not the result of model misspecification or measurement noise. Our second aim was therefore to use longitudinal data present in our large EEG databases to establish the relatively longer term (>1 year) stability of the prediction error.

As a final aim, we investigated the genetic etiology of the EEG-based predicted-age. If the prediction error is stable (that is, some subjects show systematic over- or underestimation of calendar age), and if this is to be predictive of behavioral outcomes that are known to be heritable—such as the reaching of cognitive milestones or neurodevelopmental disorders like ADHD—then the age-prediction error should also be heritable (Derks et al., 2008; de Geus, 2010; Shaw et al., 2006; Smit, de Geus, Boersma, Boomsma, & Stam, 2016). Our final aim was therefore to assess the heritability of age-prediction error by using longitudinal twin EEG datasets of children and adolescents (Boomsma, Busjahn, & Peltonen, 2002).

## Method

### Participants and procedure

In this study, two large developmental twin-family datasets with EEG recordings were used (see supplementary Table 2 for details). The first dataset was from the Netherlands Twin Register (NTR; N=836) collected as part of a study into the genetics of brain development and cognition (Boomsma et al., 2002). The EEG recordings were obtained in four waves divided into two groups. The first group of participants was measured at age 5 and 7 and a different group of participants at ages 16 and 18 (van Beijsterveldt et al., 2013). The second dataset consisted of participants taking part in a longitudinal study of Genetics, Neurocognition and Adolescence Substance Abuse (GNASA) of Washington University in St. Louis (WUSTL), and contained measurement waves at 12, 14, and 16 years (N=621).

**Table 1.**
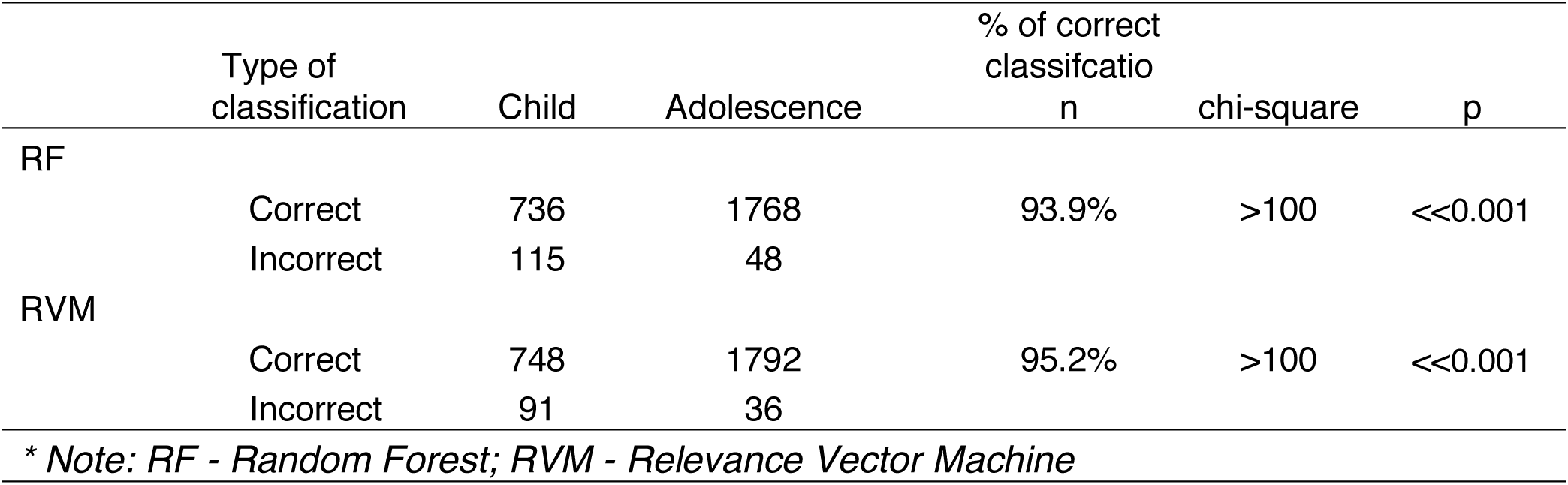
Classification childhood vs. adolescence

**Table 2.**
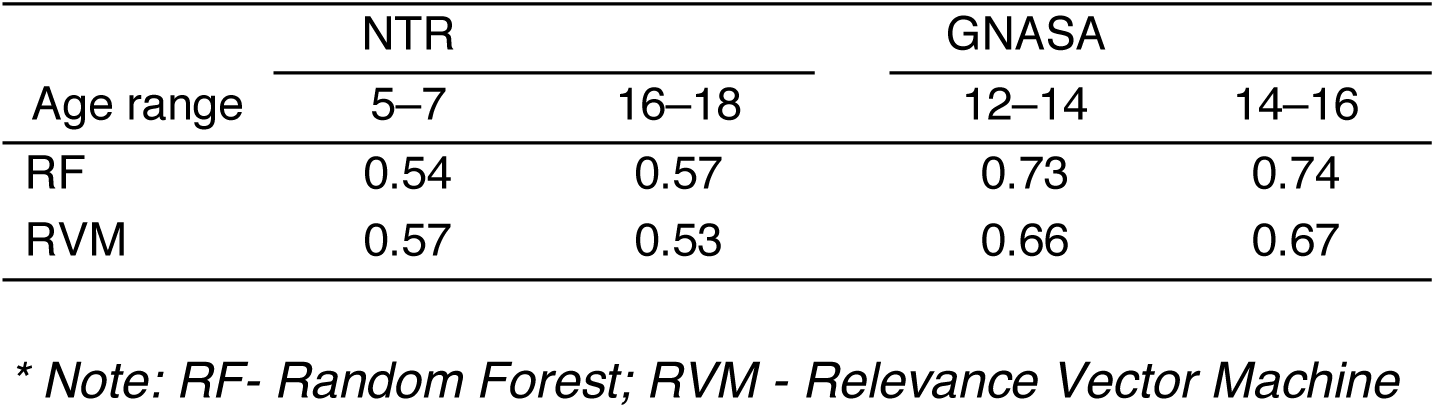
Stability of age prediction error

|  | NTR |  | GNASA |  |
| --- | --- | --- | --- | --- |
| Age range | 5–7 | 16–18 | 12–14 | 14–16 |
| RF | 0.54 | 0.57 | 0.73 | 0.74 |
| RVM | 0.57 | 0.53 | 0.66 | 0.67 |

*\* Note: RF- Random Forest; RVM - Relevance Vector Machine*

**Table 3.** Heritability (h2) of age prediction error

| Age in years | h<sup>2</sup> | p |
| --- | --- | --- |
| NTR |  |  |
| 5 | 50.8% | <0.001 |
| 7 | 62.8% | <0.001 |
| 16 | 52.8% | <0.001 |
| 18 | 42.7% | <0.001 |
| GNASA |  |  |
| 12 | 78.9% | <0.001 |
| 14 | 78.8% | <0.001 |
| 16 | 77.5% | <0.001 |

Note. Heritability was estimated in structural equation models with additive genetic (A) and Unique Environmental effects (E). Age and sex were used as covariates. Significance was determined by likelihood ratio test with 1 df.

Ethical permission for the NTR study was obtained from Medisch Ethische Toetsingscommissie (METc) of the VUmc. All participants (and parents/guardians for participants under 18) were informed about the nature of the study and were invited by letter to participate. Agreement to participate was obtained in writing. The GNASA study was approved by the human studies committee at the Washington University School of Medicine. A written informed assent was obtained from all participants, and a written informed consent was obtained from their parents.

### EEG acquisition

A detailed procedure of NTR EEG data recording is described elsewhere (Van Baal, de Geus, & Boomsma, 1996; Smit, Posthuma, Boomsma, & Geus, 2005). NTR EEG data were recorded with tin electrodes placed according to 14 channels of the 10-20 system and connected to a Nihon Kohden PV-441A polygraph with time constant 5 s (corresponding to a 0.03Hz high-pass filter) and low-pass of 35 Hz, digitized at 250 Hz using an in-house built 12-bit A/D converter board and stored for offline analysis. Leads were Fp1, Fp2, F7, F3, F4, F8, C3, C4, T5, P3, P4, T6, O1, O2, and bipolar horizontal and vertical EOG derivations. Electrode impedance was kept below 10 kΩ. All EEG signals were measured against physically connected earlobe electrodes with high impedance pre-amplifiers following PIVIK et al., (1993). Participants were seated in a dimly lit and sound attenuated booth for recording. They were instructed to close their eyes. Acquisition lasted for three periods of 1 minute. Between recording epochs, the door was opened and participants were checked not to have fallen asleep. Acquisition was extended when data was observed to have excessive artefact or sleep EEG.

The GNASA sample EEG data were recorded using Compumedics-Neuroscan SynAmps2 system from 30 scalp locations according to the extended 10-20 system using an elastic cap with Ag/AgCl electrodes and a ground electrode on the forehead, with high- and low-pass filters set at 0.05 and 100 Hz respectively on a Neuroscan SynAmps recording system. The left mastoid served as a reference during recording, and the right mastoid was recorded as a separate channel. Averaged mastoid reference was computed offline. Participants were recorded for one-minute periods with alternating eyes closed and eyes open for a total of four minutes. The data for eyes-closed were extracted from the continuous recordings.

### EEG processing

In the present study, a set of 12 channels overlapping in both NTR and WUSTL datasets was used: F3/4/7/8, C3/4, P3/4/7/8, and O1/2. All EEG signals were filtered between 1-30 Hz, individually inspected, and periods with artifacts were removed. Channels were excluded if artifact removal reduced the length of the channel signal to below the minimum total length of 90 s. EOG artifacts were removed using ICA (Delorme & Makeig, 2004), and the cleaned EEG data were partitioned into 2s epochs. Next, the signals were converted from the time domain into the frequency domain using Fast Fourier Transformation (FFT). The resulting power spectrum was divided into bins of 1 Hz, ranging from 1-24 Hz (24 bins).

GNASA and NTR samples used different apparatus and acquisition parameters (specifically, the use of different time-constants and hardware lowpass filters during recording), which could lead to age-correlated differences in power scores which could be capitalized by the machine learning algorithms for classification. We expected these spurious effects to be minimal, as the largest age range recordings (NTR childhood and adolescents, age range 5 to 18 years) are the most informative for the machine learning algorithms, and these recordings were made on the same apparatus. Nevertheless, we removed any remaining apparatus effects by removing power differences between the GNASA and NTR. Since apparatus was also confounded with age, we decided to find age-matched pairs of individuals in the GNASA and WUSTL datasets (N=30 each, Mage=16.3). PSDs were obtained for each channel, and the difference obtained between GNASA and WUSTL. These differences were then averaged across channels, since effects in apparatus were not expected to change between channels with the same settings. The averaged difference was used to correct the WUSTL power values for each 1 Hz power bin. The results were not critically affected by the removal of the apparatus/cohort effect, however, MAE increased by approximately 0.10 years. Figure S1 shows the average power spectra for these participants (NTR vs GNASA). The corrected power spectral densities for representative leads (F3 and O2) and for each wave are provided in supplementary Figure S2. **All subsequent analyses used the corrected power spectra.**

### Machine learning analyses

To estimate brain maturational level, the three most common machine learning algorithms in previous studies (see Table S1) were applied; Random Forest, Support Vector Machine and Relevance Vector Machine using power in 1 Hz wide bins from 1 to 24 Hz, corrected for sex differences in each bin, as input features to the machine learning models.

#### Random Forest

Random Forest is one of the most popular machine learning methods for classification and regression (Breiman, 2001). Random Forest creates a large number of decision trees using various bootstrapped subsamples of the data and features, a so-called random forest. To classify a new data vector based on attributes, each tree gives a classification and the tree ‘votes’ for that class. The forest chooses the classification having the most votes.

The regression extension of Random Forest works similarly, but additionally assigns a value for the outcome variable whenever a decision falls below or under a certain threshold. Random Forest improves the predictive accuracy over standard regression in cross-validation, controls for overfitting, and naturally allows for interactions between the features. In this study, the number of trees was fixed at 500 trees. The number of predictors was set to 40. Neither parameter was critical for the outcome.

#### Relevance Vector Machine (RVM)

Relevance Vector Machine is an extension of Support Vector Machine (Tipping, 2000; Vapnik, 1998). In classification Support Vector Machine, the feature data points are projected on a high-dimensional space. The classification groups are then separated using a hyperplane (Vapnik, 1998). The hyperplane is formed by the so-called support vectors, which are data points that are highly informative on the separation between classes, and close to the decision boundary. This supervised method aims to find the hyperplane that provides the largest margin between data points in the support vectors that fall into different classes. Support Vector Machine has become an important classifier over recent years.

Relevance Vector Machine utilizes a Bayesian approach to increase sparseness in the prediction. The approach aids machine learning with highly correlated predictors (as is likely to be the case in EEG data.) Relevance Vector Machine uses a probabilistic measure to define the separation space; it imposes an explicit zero-mean Gaussian prior. The relevance vectors are formed by samples appearing to be more representative of the classes, which are located away from the decision boundary of the classifier, whereas Support Vector Machine typically uses the samples close to the decision boundary as so-called support vectors. The key difference in Relevance Vector Machine compared to Support Vector Machine is that a separate hyperparameter is introduced for each of the parameters, instead of a single shared hyperparameter. When the evidence concerning these hyperparameters is maximized, a significant proportion of them go to infinity and play no role in the prediction of the model (Bishop, 2006). Therefore, the Relevance Vector Machine is a sparse classifier as the decision function depends on fewer input data that a comparable Support Vector Machine. This sparsity may lead to a faster performance on training data and results of more generalizable results by decreasing the overfitting, which can reduce error during cross-validation.

#### Validation, classification and accuracy measures

The performance of the methods was assessed in different ways. All prediction performance measures were estimated using 6-fold cross-validation. In cross-validation, all data is iteratively split into a training- and a testing set. Because of the complex familial (twin) dependence in the input datasets, we first reduced the dataset by selecting a single person from each family to avoid family relations between participants in the training and testing datasets. Next, 6-fold cross-validation was applied on the reduced set with each time the regression Relevance Vector Machine and Random Forest performed on the test set. Note that we did not use the out-of-bag option for Random Forest to maximize comparability with the Relevance Vector Machine approach. We then repeated this procedure 12 times for a different set of family members, again selecting only a single person from each family. Finally, all available prediction values were averaged.

Prediction accuracy was determined in several ways. Firstly, we used Mean Absolute Error, defined simply as the sum of the prediction errors divided by the number of recordings/measurements. Next, we assessed the wave-by-wave prediction accuracy, that is, we compared the median age to median predicted age per wave (four waves for NTR; three waves for GNASA). This method allows individual age prediction to systematically deviate from actual age without penalty, only assessing the prediction error of the wave medians compared to median actual age. Finally, we assessed longitudinal stability (correlation between waves) of the predicted age. Stability was assessed as the correlation between the prediction errors (estimated minus actual age) obtained at subsequent waves of data collection. Finally, classification accuracy was assessed for childhood versus adolescence waves of the NTR dataset.

To assess the heritability of the age-prediction errors, we used Structural Equation Modeling (SEM) implemented in R package OpenMx (Boker et al., 2011). The relative contribution of genetic and environmental effects to the total trait variance can be estimated by weighing the contribution of known levels of resemblance to the correlational structure between family members (Boomsma et al., 2002; Neale and Cardon, 1992). Specifically, additive genetic effects (A) are correlated 1 between monozygotic twin pairs (mz), and on average 0.5 between dizygotic (dz) twins and siblings. Large contributions of additive genetic effects (A), therefore, result in observed twin/sibling correlations close to these levels (r_MZ_=1, r_DZ_=0.5). Non-additive genetic effects (D)—effects of genetic dominance or epigenetic effects—are correlated 1 for MZ twins and .25 for (r_MZ_=1, r_DZ_=0.25). Common environmental effects (C)—such as the effects of rearing environment—are shared among all family members (r_MZ_=1, r_DZ_=1). Large contributions of common environmental effects (C) to the trait variance will result in high correlations equal for MZ and DZ/sibling correlations. Unique environmental effects (E) are the residual variance that cannot be explained by the familial resemblance (A or C). For twin data, unique environmental effects (E) largely reflects the variance not explained by the MZ twin correlation (1-r_MZ_). Admixtures of these effects will result in specific correlation patterns based on summing of each effect’s theoretical twin/sibling correlations weighted by the contribution of that effect. SEM iteratively searches through these relative contributions comparing the estimated and actual correlations, finishing at the maximally likely solution.

Note that in the current twin design, the contributions of C and D effects are collinear and cannot be estimated simultaneously. The Akaike Information Criterion (AIC) was used to decide which variance component (D or C) was used. We fitted ADE or ACE models to the data to estimate the relative proportions of A, C or D, and E effects on the age-prediction error of the participants. The significance of the effects was determined by fixing the estimated contribution of the effect to zero. Comparing the models with and without the effect results in a difference in likelihood of the models. Twice the difference in likelihood is approximately chi-square distributed with the reduction in free parameters as the degrees of freedom. From this p-values are obtained with the number of parameters dropped as degrees of freedom (1 df). We first tested the significance of the D or C effect. If this effect could be dropped, we established significance of heritability (A) by comparing the fit of a model with AE variance components against a model with only E.

## Results

### Machine learning prediction performance

In the full age range from childhood into adolescence, application of Random Foreset resulted in the smaller MAE of 1.22 years compared to Relevance Vector Machine yielding an MAE of 1.46 years. Both these methods were able to classify childhood vs. adolescence of NTR participants well (93.9% for Random Forest, 95.2% for Relevance Vector Machine) (see Table 1). Predicted-age wave medians were plotted against the actual age medians for all seven waves (wave age 5, 7, 16, and 18 of NTR data and ages 12, 14, and 16 of WUSTL data) (see Figure 1, red (NTR) and green (WUSTL) dots indicates the median predicted against median actual age). The figures reveal that Random Forest is not able to extrapolate beyond the minimum (4.9 years) and maximum actual age (18.5 years), resulting in bounded prediction estimates as evidenced from the very small lower error bar at age 5 and upper error bar at age 18. This phenomenon may have reduced the prediction error for Random Forest compared to Relevance Vector Machine, therefore, the lower MAE for Random Forest may not reflect real prediction accuracy,

**Figure 1.**
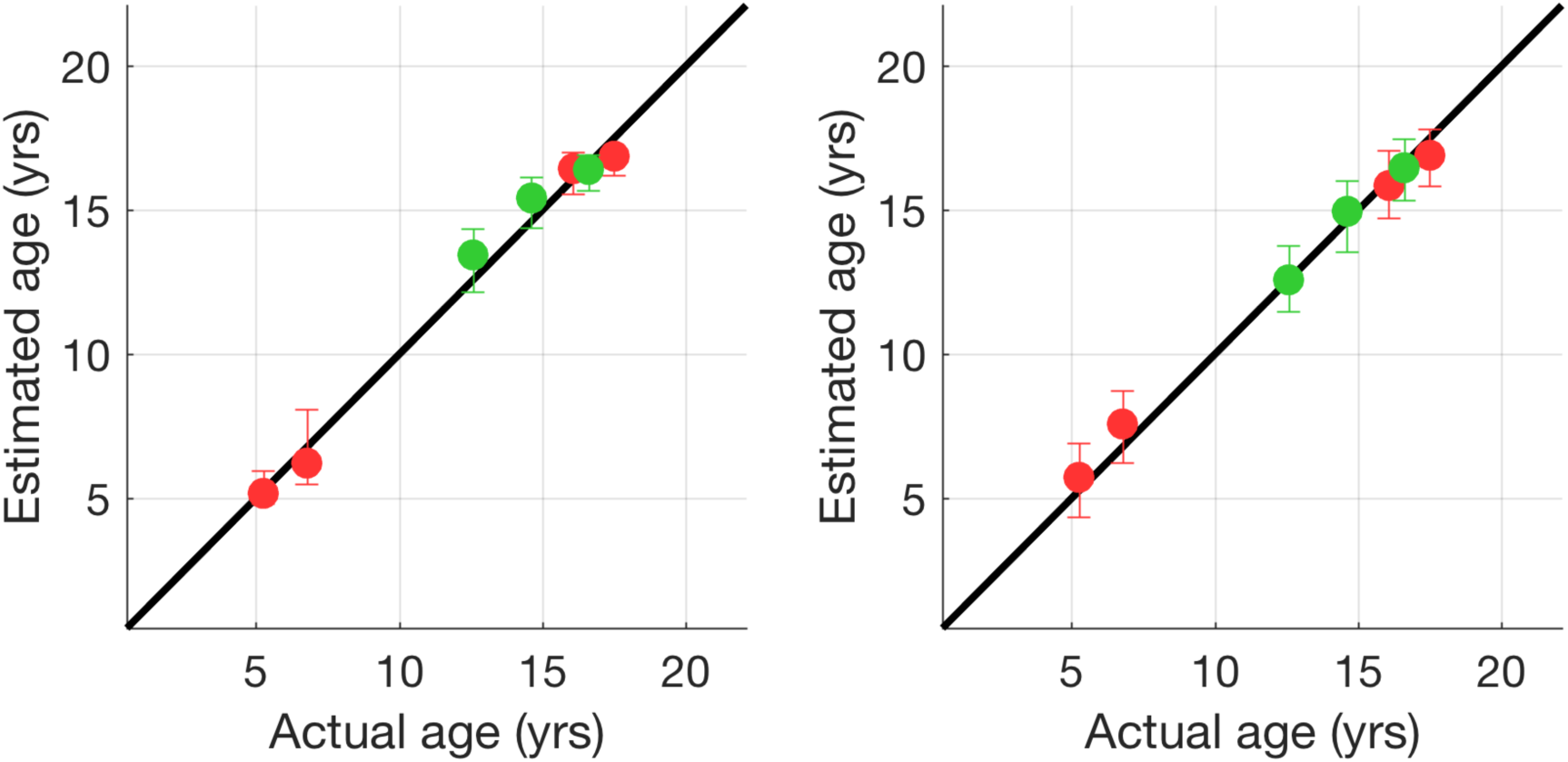
Predicted age medians plotted against actual age medians for each of the seven age groups (NTR; age 5 (n = 401,) 7 (n = 383), 16 (n = 426) and 18 (n = 368); GNASA; age 12 (n = 343), 14 (n = 425) and 16 (n = 294) of the Random Forest (right) and Relevance Vector Machine (left) algorithms. Error bars represent P75 and P25 quartiles. Wave centroids in red for NTR and green for GNASA.

A better criterion for prediction accuracy may be to look at wave centroids (i.e., average prediction age compared to average age within a wave of subjects). These wave centroids showed more error in Random Forest (average absolute error=0.503) than in Relevance Vector Machine (average absolute error=0.363). The latter showed almost perfect overlap with the perfect prediction line.

### Stability of age prediction error

For each participant, we calculated the age-prediction error as the deviation between Relevance Vector Machine or Random Forest Random Forest estimation and actual age (see Table 2). **We then correlated these between consecutive time-points across longitudinal measurements**. Prediction error stability across time was moderate to high (Random Forest: 0.54 < r < 0.74; Relevance Vector Machine Relevance Vector Machine: 0.53 < r < 0.67).

### Heritability of age prediction error

Age prediction errors from the Relevance Vector Machine predictions were entered into univariate Structural Equation Models with age and sex covariates. Common environmental (C) and nonadditive/dominant genetic effects (D) were not significant for any of the models (p>0.080), and were subsequently dropped. The best fitting models fitted only additive genetic (A) and unique environmental (E) effects. Heritability (h^2^) is then defined as the proportion of variance explained by A to the total variance (A+E). For most waves, h^2^ was substantial (h^2^>50%). For NTR age 18 the heritability was moderate (h^2^=42.7%). All heritabilities were highly significant (p<0.001).

### Contributing features in Random Forest

In order to investigate the contribution of each feature to the predictive model, we performed analysis of feature importance using the Random Forest regression only, because the random feature selection during each regression tree allows each feature to obtain a feature importance score. The lack of randomizing feature in Relevance Vector Machine and the high collinearity will obscure the importance of some features in Relevance Vector Machine, even if they are nearly as good in predicting age as the highly collinear power values at nearby electrodes and/or nearby frequencies.

Figure 2 shows the log-transformed feature importance averaged across different brain regions. The most contributing frequencies were identified at lower frequencies (delta). Other contributing frequencies were lower alpha, which changed in power but also reflected increased alpha peak frequency (see Figure S2). In addition, beta frequency power contributed to age predictions. For these frequencies, topographic plots of the feature importance from Random Forest models for these frequencies are shown in Figure 3. These indicate that the best regression was obtained with central delta, frontal lower alpha, parietal alpha, frontal lower beta, and occipital upper beta. These observations are largely consistent with the known developmental patterns in these regions (Bresnahan, Anderson, & Barry, 1999;M. & Petersen, 1973).

**Figure 2.**
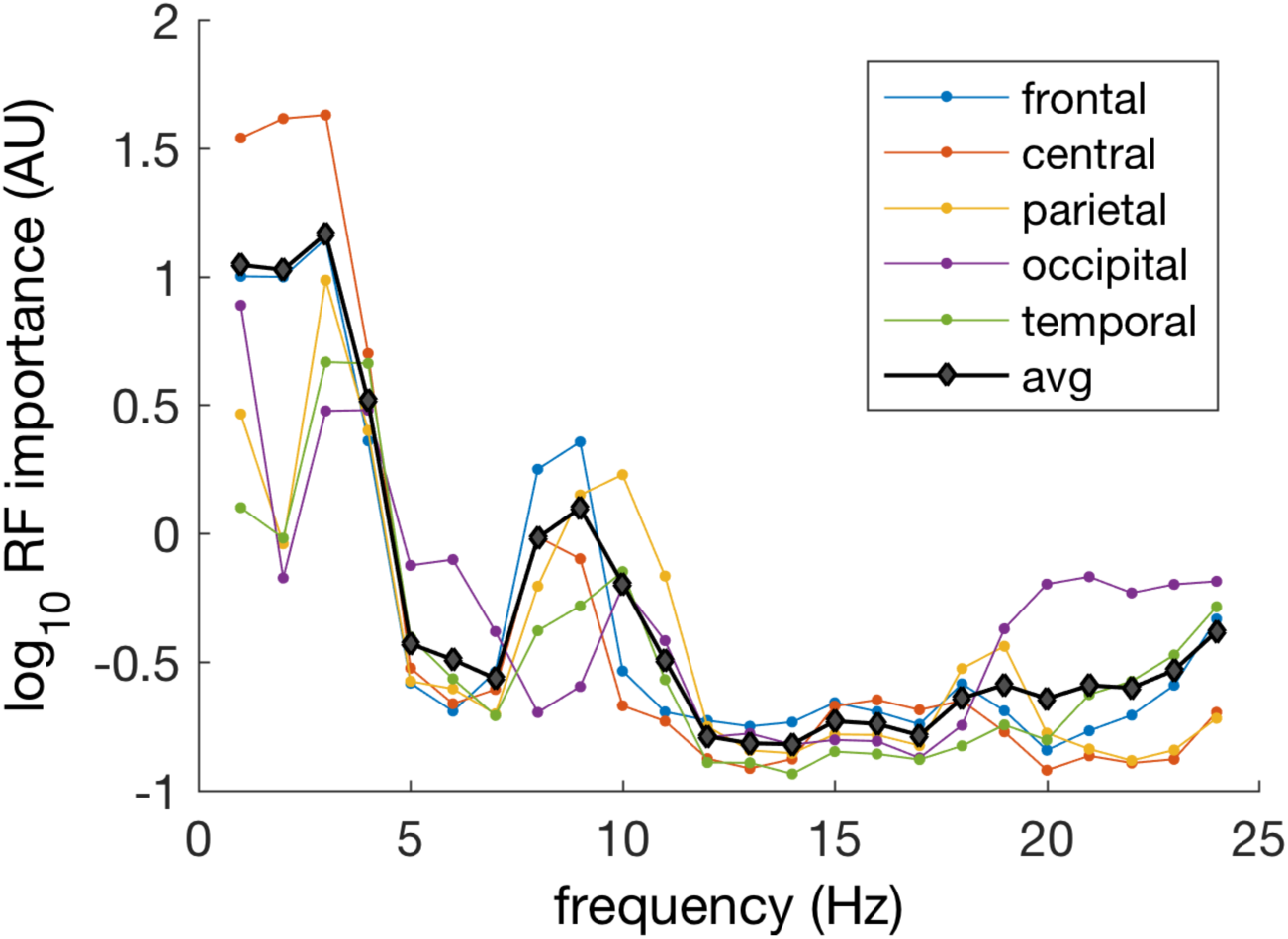
The feature importance scores were averaged across the Random Forest runs after log-transformation. The feature importance scores were then averaged for different brain regions, grand average in black.

**Figure 3.**
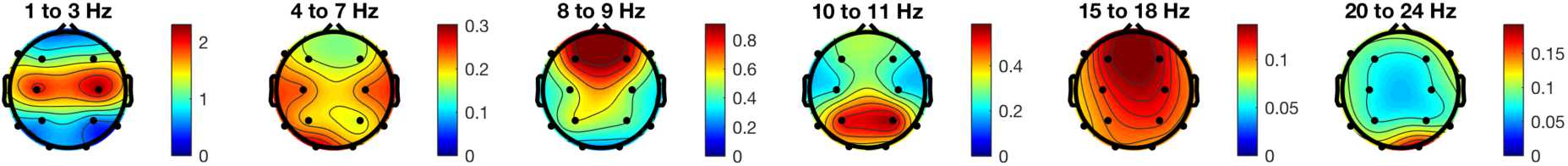
Topographic plots for the log-transformed Random Forest feature importance at the standard frequencies (delta 1 – 3 Hz, theta 4 – 7 Hz, lower alpha 8 – 9 Hz, upper alpha 10 – 11 Hz, lower beta 14 – 18 Hz, upper beta 20 – 25 Hz). The plots reveal that central delta, temporal theta, frontal lower-alpha, parietal alpha, frontal lower-beta, and occipital upper-beta are the most informative features for age classification. Note, however, that feature importance is plotted in relative strength. Low frequency power was by far the best feature for the Random Forest regression trees.

## Discussion

The results showed that brain maturational level can be estimated based on three minutes resting-state EEG recordings with high accuracy. Comparison of our outcomes based on EEG with age estimation from MRI revealed comparable results. Childhood vs. adolescent classification accuracy in MRI studies ranged from 75% - 95% (Franke, Luders, May, Wilke, & Gaser, 2012). With EEG, we obtained classification accuracy of childhood versus adolescence that was even higher (>95% for Relevance Vector Machine). The mean prediction error of 1.22 for Random Forest and 1.46 for Relevance Vector Machine are only slightly higher than the lowest prediction errors estimates obtained in MRI studies (1.1 years) in comparable age groups. The analysis of feature importance in Random Forest showed that classification depended mostly on low-frequency power; further notable contributors to the prediction were temporal theta, frontal lower alpha, parietal alpha, frontal lower beta, and occipital upper beta.

The Relevance Vector Machine and Random Forest machine learning methods were comparable in their results. One limitation of Random Forest is that it does not give fully continuous predicted ages, and cannot extrapolate outside the input age range. Random Forest prediction performance may have been artificially decreased by this capping of the output values. Relevance Vector Machine on the other hand does extrapolate, which resulted in higher mean prediction error, but also yielded illogical age estimates (even below zero for two participants), suggesting that the remaining age-prediction error has variability that cannot logically be attributed to brain maturation. On the other hand, Relevance Vector Machine did provide the best fit of age-group median centroids, outperforming Random Forest. In our view, these results combined suggest that sparser models such as the Relevance Vector Machine are superior for age prediction using EEG power, even though remaining error variance is likely to exist.

The over- or underestimation relative to actual age indicates that some children and adolescents exhibit EEG brain activity patterns more like those of younger or older ages. This study is the first to show that this prediction error was systematic, and moderately to highly stable over a period of 1.5 years (0.54 < r 0.74 for Random Forest and 0,53 < r < 0.67 for Relevance Vector Machine). In addition, age-prediction error was to a large extent heritable, ranging from 43% to 79%. No significant effects of shared environment or non-additive genetic effects—i.e. the interactions among alleles both within and across gene loci—were found, suggesting that the systematic deviation is largely genetically determined, but with substantial unique environmental influences that include non-stable error estimates. Although it has yet to be determined whether these stable and heritable deviations are predictive of behavioral traits and/or neurodevelopmental disorders, the fact that they do not fully reflect unstable variance due to measurement noise or model misspecification indicates that prediction error is a good candidate for predicting stable individual differences in neurodevelopment, cognition, and neurodevelopmental disorders.

Previous research has suggested that scoring high or low on selected behavioral phenotypes (viz., IQ and ADHD) are associated with slower or faster brain-maturation trajectories (Shaw et al., 2006, 2007, 2012; Sowell, 2004). These studies were the first to suggest that brain maturational level estimated from brain parameters is a property correlated with behavioral outcomes. These neurodevelopmental phenotypes are therefore prime candidates to link to the EEG-based predicted age, Moreover, attention problems and IQ are to a large extent genetically determined (Derks et al., 2008; Posthuma, Mulder, Boomsma, & De Geus, 2002). Since we have shown that EEG-based predicted age has a genetic etiology, this opens up the possibility for correlating the stable and heritable maturational trajectories with these genetically-mediated behavioral outcomes, and establish the genetic overlap between the traits. These correlations have yet to be established.

Overall, our results show that age predictions from low-cost EEG recordings can be performed with a precision comparable to predictions obtained from MRI in an age range from childhood to adolescence. In addition, we have shown that the prediction error is not random noise, but moderately stable over a period of about 1.5 years, and to a large extent influenced by the genetic background of the subject. These findings clear the way for EEG-based age prediction as a marker of brain maturation and investigation of its relation with (genetically mediated) neurodevelopmental phenotypes, such as cognitive performance and ADHD. In clinical practice, age prediction—and especially the systematic deviation from actual age—may prove a valuable biomarker for neuropsychiatric disorders related to abnormal brain development, or normal behavioral outcomes. In parallel with the epigenetic clock (Horvath, 2013; Jylhävä, Pedersen, & Hägg, 2017; Jones, Goodman, & Kobor, 2015), EEG-estimated brain maturation might become a tool that address questions concerning developmental trajectories. Some studies have shown a relation between brain maturational level and behavioral phenotypes (ADHD, impulsivity, and IQ; Shaw et al., 2007, 2012; Yang et al., 2015). These studies showed that ADHD and superior IQ both were related to a delay in maturation. Future studies may show that this relation can also be captured by EEG-estimated brain maturation, and could therefore be an important additional predictor of scholastic achievement and attention problems.

## Acknowledgements

This research was supported by grants BBR Foundation (NARSAD) Young Investigator grant 21668 and NWO/MagWVENI-451-08-026 to D.S.; National Institutes of Health DA018899 and DA027096 to A.P.A.

## References

Anderson, V., Northam, E., Hendy, J., & Wrennall, J. (2001). Developmental neuropsychology: a clinical approach.

Barriga-Paulino, C., Flores, A., & Gomez, C. (2011). Developmental Changes in the EEG Rhythms of Developmental Changes in the EEG Rhythms of Children and Young Adults Analyzed by Means of Correlational, Brain Topography and Principal Component Analysis. Journal of Psychophysiology, 25(3), 143–158. https://doi.org/10.1027/0269-8803/a000052

Benniger, C., Matthis, P., & Scheffner, D. (1984). EEG development of healthy boys and girls: results of a longitudinal study. Electroencephalography and Clinical Neurophysiology, 57(1), 1–12.

Bishop, C. M. (2006). Pattern Recognition and Machine Learning. Journal of Chemical Information and Modeling (Vol. 53). https://doi.org/10.1117/1.2819119

Boker, S., Neale, M., Maes, H., Wilde, M., Spiegel, M., Brick, T., … Fox, J. (2011). OpenMx: An Open Source Extended Structural Equation Modeling Framework. Psychometrika, 76(2), 306–317. https://doi.org/DOI/10.1007/s11336-010-9200-6

Boomsma, D., Busjahn, A., & Peltonen, L. (2002). Classical twin studies and beyond. Nature Reviews Genetics, 3(11), 872–882. https://doi.org/10.1038/nrg932

Boomsma, D. I., Vink, J. M., van Beijsterveldt, T. C., de Geus, E. J., Beem, A. L., Mulder, E. J., … van Baal, G. C. (2002). Netherlands Twin Register: a focus on longitudinal research. Twin Research, 5(5), 401–406. https://doi.org/10.1375/136905202320906174

Breiman, L. (2001). Random Forests. Machine Learning, 1–33.

Bresnahan, S. M., Anderson, J. W., & Barry, R. J. (1999). Age-Related Changes in Quantitative EEG in Attention-Deficit / Hyperactivity Disorder.

Brown, T. T., Kuperman, J. M., Chung, Y., Erhart, M., McCabe, C., Hagler, D. J., … Dale, A. M. (2012). Neuroanatomical assessment of biological maturity. Current Biology, 22(18), 1693–1698. https://doi.org/10.1016/j.cub.2012.07.002

de Geus, E. J. C. (2010). From genotype to EEG endophenotype: a route for post-genomic understanding of complex psychiatric disease? GENOME MEDICINE, 2. https://doi.org/10.1186/gm184

Delorme, A., & Makeig, S. (2004). EEGLAB: an open source toolbox for analysis of single-trial EEG dynamics including independent component analysis. Journal of Neurscience Methods, 134, 9–21. https://doi.org/10.1016/j.virol.2015.08.001

Derks, E. M., Hudziak, J. J., Dolan, C. V., Van Beijsterveldt, T. C. E. M., Verhulst, F. C., & Boomsma, D. I. (2008). Genetic and environmental influences on the relation between attention problems and attention deficit hyperactivity disorder. Behavior Genetics, 38(1), 11–23. https://doi.org/10.1007/s10519-007-9178-8

Durston, S., Casey, B., Durston, S., & Casey, B. J. (2006). What have we learned about cognitive development from neuroimaging ?, (February). https://doi.org/10.1016/j.neuropsychologia.2005.10.010

Franke, K., Luders, E., May, A., Wilke, M., & Gaser, C. (2012). Brain maturation: Predicting individual BrainAGE in children and adolescents using structural MRI. NeuroImage, 63(3), 1305–1312. https://doi.org/10.1016/j.neuroimage.2012.08.001

Franke, K., Ziegler, G., Klöppel, S., & Gaser, C. (2010). Estimating the age of healthy subjects from T1-weighted MRI scans using kernel methods: Exploring the influence of various parameters. NeuroImage, 50(3), 883–892. https://doi.org/10.1016/j.neuroimage.2010.01.005

Gasser, T., Jennen-Steinmetz, C., Sroka, L., Verleger, R., & Mocks, J. (1988). Development of the EEG of school-age children and adolescents. Electroencephalography and Clinical Neurophysiology, 69(2), 100–109.

Gasser, T., Verleger, R., Bächer, P., & Sroka, L. (1988). Development of the EEG of school-age children and adolescents. I. Analysis of band power. Electroencephalography and Clinical Neurophysiology, 69(2), 91–99. https://doi.org/10.1016/0013-4694(88)90204-0

Giedd, J. N., Lalonde, F. M., Celano, M. J., White, S. L., Wallace, G. L., Lee, N. R., & Lenroot, R. K. (2009). Anatomical Brain Magnetic Resonance Imaging of Typically Developing Children and Adolescents. Journal of the American Academy of Child & Adolescent Psychiatry, 48(5), 465–470. https://doi.org/10.1097/CHI.0b013e31819f2715

Hedman, A. M., van Haren, N. E. M., Schnack, H. G., Kahn, R. S., & Hulshoff Pol, H. E. (2012). Human brain changes across the life span: A review of 56 longitudinal magnetic resonance imaging studies. Human Brain Mapping, 33(8), 1987–2002. https://doi.org/10.1002/hbm.21334

Horvath, S. (2013). DNA methylation age of human tissues and cell types. Genome Biol, 14(10), R115. https://doi.org/10.1186/gb-2013-14-10-r115

Huttenlocher, P. R. (1979). Synaptic density in human frontal cortex - Developmental changes and effects of aging. Brain Research, 163(2), 195–205. https://doi.org/10.1016/0006-8993(79)90349-4

Jones, M. J., Goodman, S. J., & Kobor, M. S. (2015). DNA methylation and healthy human aging. Aging Cell, 14(6), 924–932. https://doi.org/10.1111/acel.12349

Jylhävä, J., Pedersen, N. L., & Hägg, S. (2017). Biological Age Predictors. EBioMedicine, 21, 29–36. https://doi.org/10.1016/j.ebiom.2017.03.046

Lenroot, R. K., & Giedd, J. N. (2006). Brain development in children and adolescents : Insights from anatomical magnetic resonance imaging, 30, 718–729. https://doi.org/10.1016/j.neubiorev.2006.06.001

Matousek, M., & Petersen, I. (1973a). Frequency analysis of the EEG in normal children and adolescents. In Automation of Clinical Electroencephalography (pp. 75–102).

Matousek, M., & Petersen, I. (1973b). Frequency analysis of the EEG in normal children and normal adolescents. Automation of Clinical Electroencephalography, 75–102.

Mills, K. L., & Tamnes, C. K. (2014). Developmental Cognitive Neuroscience Methods and considerations for longitudinal structural brain imaging analysis across development. Accident Analysis and Prevention, 9, 172–190. https://doi.org/10.1016/j.dcn.2014.04.004

Niedermeyer, E., & Lopes Da Silva, F. (2005). Electroencephalography: basic principles, clinical applications, and related fiels. lippincott Williams & Wilkins.

Pivik, R. T., Broughton, R. J., Coppola, R., Davidson, R. J., Fox, N., & Nuwer, M. R. (1993). Guidelines for the recording and quantitative analysis of electroencephalographic activity in research contexts. Psychophysiology. https://doi.org/10.1111/j.1469-8986.1993.tb02081.x

Posthuma, D., Mulder, E. J. C. M., Boomsma, D. I., & De Geus, E. J. C. (2002). Genetic analysis of IQ, processing speed and stimulus-response incongruency effects. Biological Psychology, 61(1–2), 157–182. https://doi.org/10.1016/S0301-0511(02)00057-1

Puligheddu, M., De Munck, J. C., Stam, C. J., Verbunt, J., De Jongh, A., Van Dijk, B. W., & Marrosu, F. (2005). Age distribution of MEG spontaneous theta activity in healthy subjects. Brain Topography, 17(3), 165–175. https://doi.org/10.1007/s10548-005-4449-2

Shaw, P., Eckstrand, K., Sharp, W., Blumenthal, J., Lerch, J. P., Greenstein, D., … Rapoport, J. L. (2007). Attention-deficit/hyperactivity disorder is characterized by a delay in cortical maturation. Proceedings of the National Academy of Sciences, 104(49), 19649–19654. https://doi.org/10.1073/pnas.0707741104

Shaw, P., Greenstein, D., Lerch, J., Clasen, L., Lenroot, R., Gogtay, N., … Giedd, J. (2006). Intellectual ability and cortical development in children and adolescents. Nature, 440(7084), 676–679. https://doi.org/10.1038/nature04513

Shaw, P., Lerch, J., Greenstein, D., Sharp, W., Clasen, L., Evans, A., … Rapoport, J. (2006). Longitudinal Mapping of Cortical Thickness and Clinical Outcome in Children and Adolescents With Attention-Deficit/Hyperactivity Disorder. Archives of General Psychiatry, 63(May 2006), 540–549. https://doi.org/10.1001/archpsyc.63.5.540.ABSTRACT

Shaw, P., Malek, M., Watson, B., Sharp, W., Evans, A., & Greenstein, D. (2012). Development of cortical surface area and gyrification in attention-deficit/hyperactivity disorder. Biological Psychiatry, 72(3), 191–197. https://doi.org/10.1016/j.biopsych.2012.01.031

Smit, D. J. A., E. J. C. De Geus, Boersma, M., Boomsma, D. I., & Stam, C. J. (2016). Life-Span Development of Brain Network, 6(4), 312–325. https://doi.org/10.1089/brain.2015.0359

Smit, D. J. A., Posthuma, D., Boomsma, D. I., & Geus, E. J. C. D. E. (2005). Heritability of background EEG across the power spectrum. Psychophysiology, 42, 691–697. https://doi.org/10.1111/j.1469-8986.2005.00352.x

Sowell, E. R. (2004). Longitudinal Mapping of Cortical Thickness and Brain Growth in Normal Children. Journal of Neuroscience, 24(38), 8223–8231. https://doi.org/10.1523/JNEUROSCI.1798-04.2004

Tipping, M. E. (2000). The Relevance Vector Machine. Advances inNeural InformationProcessingSystems 12. https://doi.org/10.1.1.34.4986

Van Baal, G. C. M., de Geus, E. J. C., & Boomsma, D. I. (1996). Genetic architecture of EEG power spectra in early life. Electroencephalography and Clinical Neurophysiology, 98, 502–514.

van Beijsterveldt, C. E. M., Groen-Blokhuis, M., Hottenga, J. J., Franić, S., Hudziak, J. J., Lamb, D., … Boomsma, D. I. (2013). The Young Netherlands Twin Register (YNTR): longitudinal twin and family studies in over 70,000 children. Twin Research and Human Genetics : The Official Journal of the International Society for Twin Studies, 16(1), 252–67. https://doi.org/10.1017/thg.2012.118

Vapnik, V. N. (1998). Statistical Learning Theory. New York: Wiley. https://doi.org/10.2307/1271368

Walhovd, K. B., Tamnes, C. K., & Fjell, A. M. (2014). Brain structural maturation and the foundations of cognitive behavioral development, 27(2), 176–184. https://doi.org/10.1097/WCO.0000000000000074

Yang, Y., Wang, P., Baker, L. A., Narr, K. L., Joshi, S. H., Hafzalla, G., … Thompson, P. M. (2015). Thicker temporal cortex associates with a developmental trajectory for psychopathic traits in adolescents. PLoS ONE, 10(5), 1–15. https://doi.org/10.1371/journal.pone.0127025

